# Lactate dehydrogenase A-coupled NAD^+^ regeneration is critical for acute myeloid leukemia cell survival

**DOI:** 10.1101/2025.03.03.641255

**Authors:** Ayşegül Erdem, Séléna Kaye, Francesco Caligiore, Manuel Johanns, Fleur Leguay, Jan Jacob Schuringa, Keisuke Ito, Guido Bommer, Nick van Gastel

## Abstract

**Background:** Enhanced glycolysis plays a pivotal role in fueling the aberrant proliferation, survival and therapy resistance of acute myeloid leukemia (AML) cells. Here, we aimed to elucidate the extent of glycolysis dependence in AML by focusing on the role of lactate dehydrogenase A (LDHA), a key glycolytic enzyme converting pyruvate to lactate coupled with the recycling of NAD^+^.

**Methods:** We compared the glycolytic activity of primary AML patient samples to protein levels of metabolic enzymes involved in central carbon metabolism including glycolysis, glutaminolysis and the tricarboxylic acid cycle. To evaluate the therapeutic potential of targeting glycolysis in AML, we treated AML primary patient samples and cell lines with pharmacological inhibitors of LDHA and monitored cell viability. Glycolytic activity and mitochondrial oxygen consumption were analyzed in AML patient samples and cell lines post-LDHA inhibition. Perturbations in global metabolite levels and redox balance upon LDHA inhibition in AML cells were determined by mass spectrometry, and ROS levels were measured by flow cytometry.

**Results:** Among metabolic enzymes, we found that LDHA protein levels had the strongest positive correlation with glycolysis in AML patient cells. Blocking LDHA activity resulted in a strong growth inhibition and cell death induction in AML cell lines and primary patient samples, while healthy hematopoietic stem and progenitor cells remained unaffected. Investigation of the underlying mechanisms showed that LDHA inhibition reduces glycolytic activity, lowers levels of glycolytic intermediates, decreases the cellular NAD^+^ pool, boosts OXPHOS activity and increases ROS levels. This increase in ROS levels was however not linked to the observed AML cell death. Instead, we found that LDHA is essential to maintain a correct NAD^+^/NADH ratio in AML cells. Continuous intracellular NAD^+^ supplementation via overexpression of water-forming NADH oxidase from *Lactobacillus brevis* in AML cells effectively increased viable cell counts and prevented cell death upon LDHA inhibition.

**Conclusions:** Collectively, our results demonstrate that AML cells critically depend on LDHA to maintain an adequate NAD^+^/NADH balance in support of their abnormal glycolytic activity and biosynthetic demands, which cannot be compensated for by other cellular NAD^+^ recycling systems. These findings also highlight LDHA inhibition as a promising metabolic strategy to eradicate leukemic cells.

## Background

Acute myeloid leukemia (AML) is a heterogeneous hematological malignancy characterized by the rapid growth of abnormal myeloid progenitor cells in the bone marrow, leading to impaired hematopoiesis [1, 2]. Despite therapeutic advances, patients continue to experience poor outcomes and high rates of relapse, highlighting the need for a more profound understanding of the molecular mechanisms that drive leukemogenesis.

Since the initial observations of Otto Warburg regarding the reliance of cancer cells on aerobic fermentation for their energy production [3], extensive research over more then a century has underscored the importance of metabolic abnormalities in cancer development. Yet, the complex relationship between disrupted cellular metabolism and oncogenic progression has only started to become elucidated. Emerging evidence indicates that dysregulated metabolism not only fuels the aberrant proliferation and survival of AML tumors [4], but also plays a pivotal role in chemoresistance [5–9]. While metabolic profiles can differ significantly between AML patients and even within AML tumors [10–12], glycolysis has been shown to be essential for AML blasts in most patient populations, distinguishing itself from other metabolic pathways by providing rapid ATP production, crucial biosynthetic precursors, and an adaptive advantage in the hypoxic bone marrow microenvironment [13, 14]. In accordance, the glycolytic metabolites lactate and pyruvate are elevated in AML patient serum samples [15]. Additionally, interactions with the bone marrow microenvironment enhance glycolysis in AML cells, with this glycolytic shift being regulated by the CXCL12/CXCR4/mTOR signaling axis, which serves as a protective mechanism against chemotherapy [16]. Pyruvate dehydrogenase kinase (PDK) enzymes, which inhibit the entrance of pyruvate into the mitochondrial tricarboxylic acid (TCA) cycle, have been shown to fine-tune glycolysis by suppressing mitochondrial oxidative phosphorylation (OXPHOS) not only in leukemic blasts [10], but also in leukemia-initiating cells (also known as leukemic stem cells; LSCs) [13]. LSCs, while generally considered to depend more on OXPHOS, thus also require a functional glycolytic pathway, and this further increases when AML cells are treated with OXPHOS inhibitors [17, 18]. Despite the overall importance of glycolysis in AML, the role of individual glycolytic enzymes has been explored much less in depth. A better characterization of the differential dependency of AML cells on specific glycolytic enzymes could offer new therapeutic opportunities.

Lactate dehydrogenase A (LDHA), a predominant enzyme in aerobic and anaerobic glycolysis, plays a pivotal role in supporting the energetic and biosynthetic demands of cancer cells. By catalyzing the conversion of pyruvate to lactate, LDHA facilitates the regeneration of nicotinamide adenine dinucleotide (NAD^+^), thereby promoting glycolytic flux and maintaining redox balance. As such, LDHA also diverts glucose-derived pyruvate away from other metabolic fates, such as entry into the mitochondrial TCA cycle for further oxidation or conversion into alanine. In the context of AML, dysregulated LDHA activity has been implicated in leukemogenesis and disease progression, highlighting its potential as a therapeutic target. A pioneering study by Wang *et al.* showed that loss of the glycolytic enzymes PKM2 (pyruvate kinase isoform M2) and LDHA has different effects on healthy and leukemic hematopoietic stem and progenitor cells (HSPCs) in mice [19]. While loss of PKM2, which lowers but does not block glycolytic flux, negatively affects hematopoietic progenitor cells (HPCs) and AML cells, hematopoietic stem cells (HSCs) are spared. Complete inhibition of fermentation by loss of LDHA in contrast equally impacts HSCs, HPCs and AML cells. However, while the effects of LDHA loss in HSCs could be ascribed to increased mitochondrial respiration and associated reactive oxygen species (ROS) generation, the role of LDHA in AML cells was independent of ROS buffering capacity and remains unknown. The importance of LDHA for human AML cells has also not yet been explored.

Considering the differential importance and regulation of glycolysis in healthy and malignant hematopoiesis, a better understanding of the specific roles of individual glycolytic enzymes is needed to design therapeutic strategies that target AML cells while maximally sparing their non-malignant counterparts. Here, we set out to explore the mechanistic interplay between LDHA-mediated metabolism and human AML pathophysiology by dissecting the direct cellular and molecular consequences of LDHA inhibition in AML primary patient samples and cell lines.

## Results

### Human AML cells are sensitive to LDHA enzyme inhibition

To better understand the role of individual enzymes in the regulation of glycolytic activity in AML cells, we re-analyzed our previously generated data [10] and correlated protein levels of central carbon metabolism enzymes (including those involved in glycolysis, the TCA cycle and glutaminolysis) of primary AML patient blasts (n=13; CD34^+^ fraction, or total mononuclear cells in case of NPM1 mutant AML) with their glycolytic activity as measured by the extracellular acidification rate (ECAR). Pearson [10] correlation analysis showed that LDHA protein levels had the highest positive correlation with ECAR activity in the patient blasts (Fig. 1A). Protein levels of pyruvate dehydrogenase phosphatase 1 (PDP1), which promotes shuttling of pyruvate towards the TCA cycle, was found to have the most negative correlation with the ECAR response (Fig. 1A). To validate this observation, we treated different human AML cell lines (n=5) and primary patient blasts (CD34^+^ cells, n=3) with a number of widely used inhibitors related to lactate/pyruvate metabolism: AZD3965 (an inhibitor of the lactate transporter monocarboxylate transporter 1; MCT1), Compound 3k (an inhibitor of PKM2, which regulates the final and rate-limiting step of glycolysis by converting phosphoenolpyruvate to pyruvate) and FX11 (an inhibitor of LDHA). We found that AML cells were highly sensitive to FX11 compared to the other two inhibitors whereby FX11 treatment also showed comparable susceptibility across different AML subtypes (Fig. 1C, Supplementary Table 1). To confirm this, we treated a range of human AML cell lines with FX11 or an unrelated selective LDHA inhibitor (GSK2837808A) and measured both viable cell numbers and the percentage of dead cells 48 hours later. In all cell lines tested, and for both LDHA inhibitors, a strong induction of cell death was observed (Fig. 1D-G). We also treated a panel of AML primary patient blasts (n=12; CD34^+^ fraction, or total mononuclear blasts in case of CD34^-^ AMLs; see Supplementary Table 1) with the two LDHA inhibitors for 48 hours and found a reduced number of viable cells upon LDHA inhibition across samples (Fig. 1H). To check possible toxic effects of LDHA inhibition on healthy HSPCs, we tested cord blood-derived CD34^+^ cells (n=3 different donors), which did not show any signs of cell death after 48 hours of treatment with 20µM FX11 (Fig. 1I, J). These data underscore the importance of LDHA for maintaining the viability of AML cells and indicate the existence of a differential dependency on this metabolic enzyme compared to healthy HSPCs.

**Figure 1.**
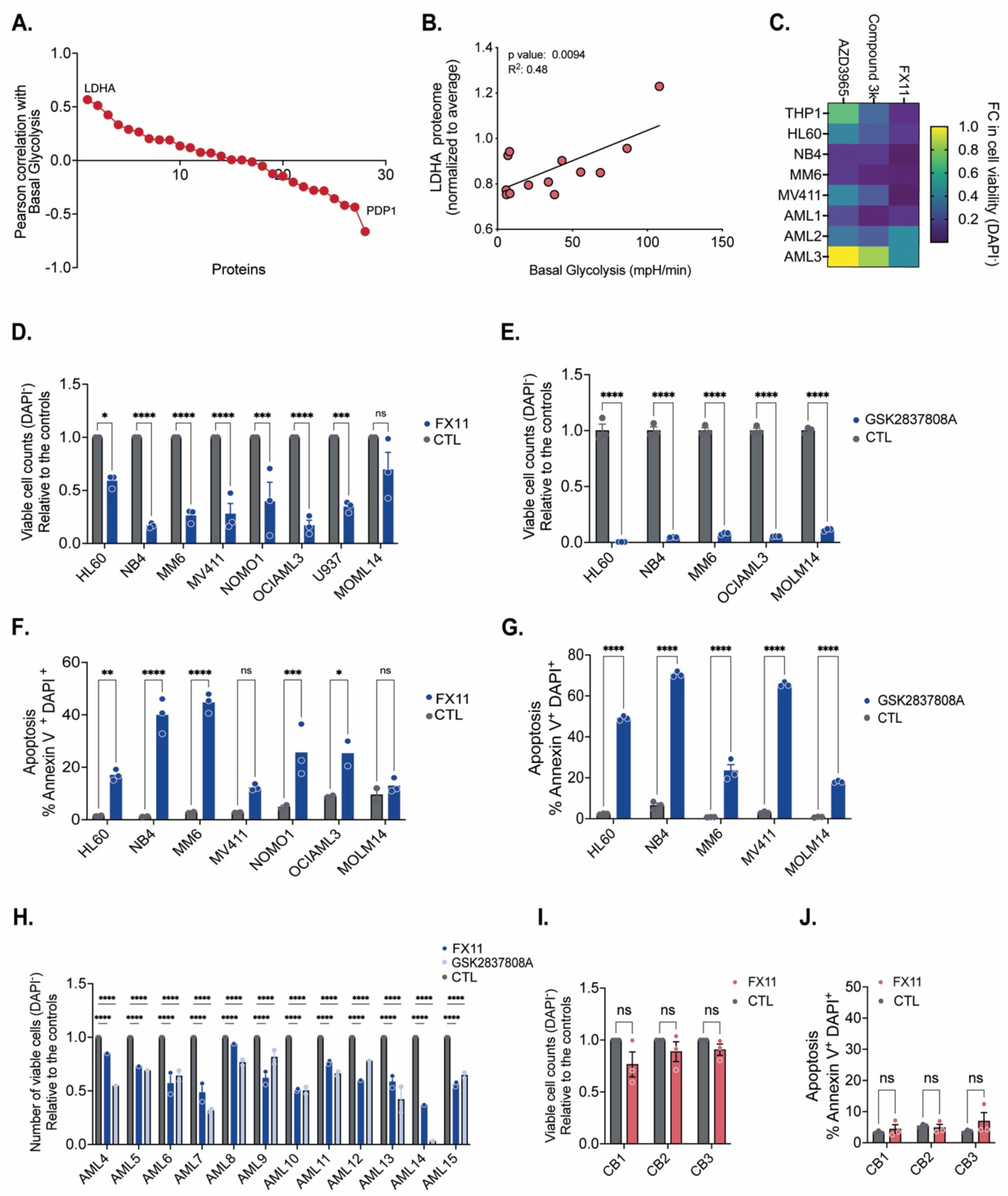
Human AML cells are sensitive to LDHA inhibition. **A.** Pearson correlation of protein levels of central carbon metabolism enzymes involved in glycolysis, the TCA cycle and glutaminolysis of primary AML patient blasts (n=13) versus their glycolytic activity as measured by the extracellular acidification rate (ECAR) (data retrieved from Erdem *et al*. [10]). **B**. Correlation of LDHA protein expression compared to ECAR activity of CD34^+^ AML primary samples (*n* = 13). **C**. Heatmap showing fold change in number of viable (DAPI^-^) AML cells (n=5 cell lines, n=3 AML primary patient samples) 24 hours after treatment with FX11 (20 µM), AZD3965 (4 µM) or Compound 3k (5 µM). **D-E**. Number of viable (DAPI^-^) cells relative to the controls in AML cells following 20 µM FX11 (D) or 20 µM GSK2837808A (E) treatment for 24 hours. Each dot represents a biological replicate measured in technical triplicate. **F-G**. Percentage dead (Annexin V^+^DAPI^+^) AML cells upon 20 µM FX11 (F) or 20 µM GSK2837808A (G) treatment for 24 hours. Each dot represents a biological replicate measured in technical triplicate. **H**. Number of viable (DAPI^-^) cells relative to controls in AML primary patient samples following 20 µM FX11 or 20 µM GSK2837808A treatment for 48 hours. Each dot represents a technical replicate. **I-J**. Number of viable (DAPI^-^) cells relative to controls (I) and percentage dead (Annexin V^+^DAPI^+^) (J) healthy cord blood-derived CD34^+^ cells following 20 µM FX11 or 20 µM GSK2837808A treatment for 48 hours. Each dot represents a biological replicate measured in technical triplicate. **B**: linear regression analysis; **D,E,F,G,I,J**: one-way ANOVA; **H**: two-way ANOVA for multiple comparisons. All experiments: lines and error bars represent mean ± SEM. \**p* < 0.05; \*\**p* < 0.01; \*\*\**p* < 0.001; \*\*\*\**p* < 0.0001; ns, not significant.

### LDHA inhibition reduces glycolysis and increases OXPHOS activity in AML cells

To investigate the metabolic consequences of LDHA inhibition, we monitored glycolysis and mitochondrial OXPHOS activity in AML cells by respectively measuring the ECAR and oxygen consumption rate (OCR) using extracellular flux analysis (Seahorse assay). Cells were first exposed to either of the two LDHA inhibitors, followed by injection of glucose, oligomycin (to inhibit mitochondrial ATP synthase) and 2-DG (2-deoxyglucose; a glucose analog and upstream inhibitor of glycolysis) into the culture medium. NB4 and HL60 cells showed a significantly lower ECAR and higher OCR after 10 minutes of LDHA inhibition, indicating a reduction in glycolytic activity (Fig. 2A-D) and a compensatory increase in OXPHOS (Fig. 2E, F). We confirmed this observation in several primary AML patient blasts (AML4, AML6, AML11; see Fig. 1H and Supplementary Table 1) that, when pre-treated with GSK2837808A, significantly reduced their ECAR and increased the OCR (Fig. 2G, H). This suggests that pharmacological LDHA inhibition reverses the Warburg effect and boosts oxidative phosphorylation in AML cells.

**Figure 2.**
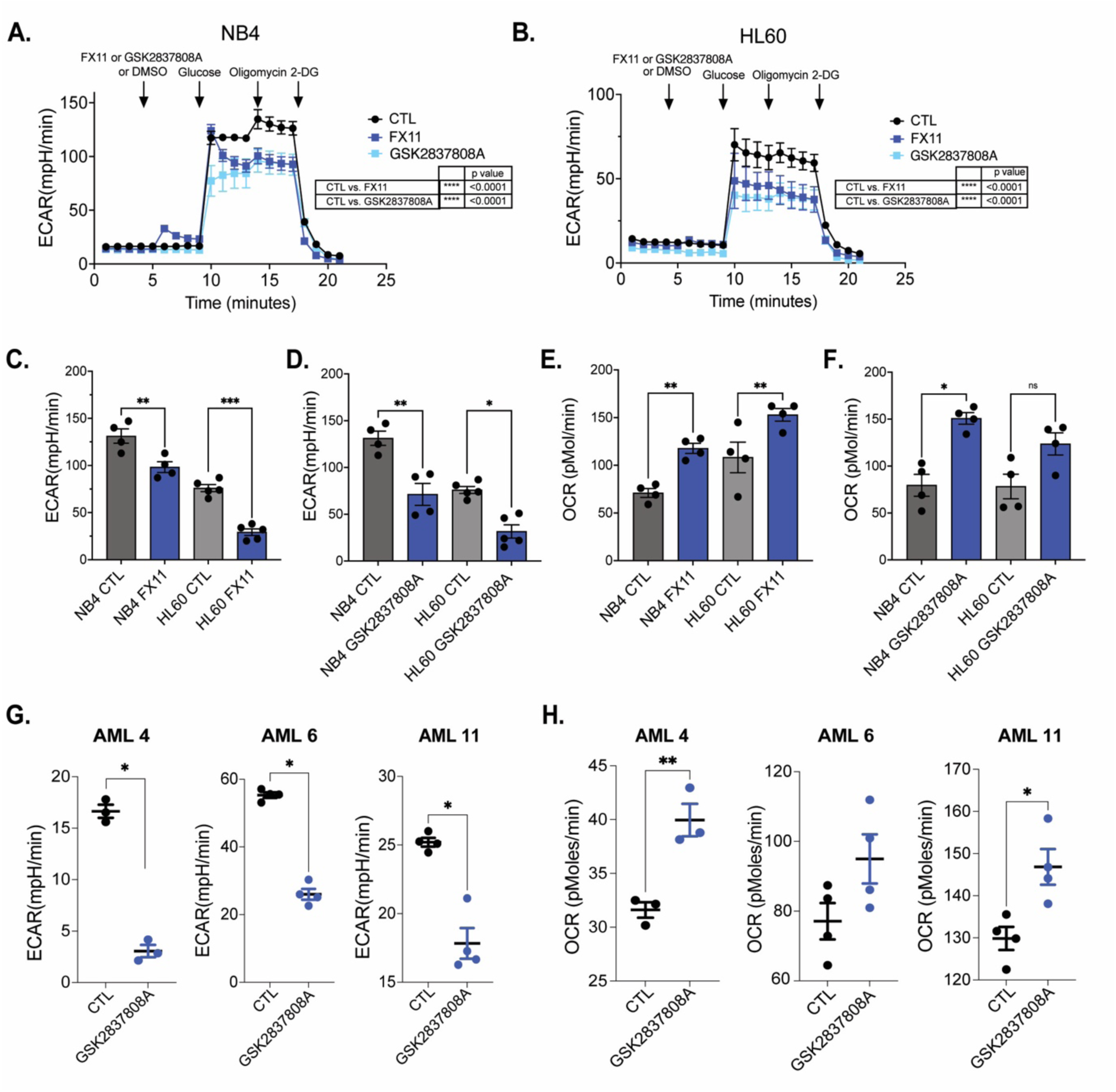
Inhibiting LDHA decreases glycolytic activity while promoting oxidative phosphorylation in AML cells. **A-B**. Real-time extracellular acidification rate (ECAR) of AML cell lines NB4 (A) and HL60 (B) (mean ± SEM of biological triplicates) measured by Seahorse bioassay after sequential injections of either of the two LDHA inhibitors (20 µM of FX11, 10 µM of GSK2837808A), glucose, oligomycin and 2-DG. **C-F**. Bar graphs showing glycolytic capacity (C: FX11, D: GSK2837808A) and maximal oxygen consumption rate (OCR; E: FX11, F: GSK2837808A) of NB4 and HL60 AML cells. **G-H**. Basal ECAR (G) and OCR (H) after pre-treatment of AML primary patient samples (n=3, each dot represents a technical replicate) with 10 µM GSK2837808A. **G**,**H**: Student’s t-test; **C,D,E,F**: one-way ANOVA; **A,B**: two-way ANOVA for multiple comparisons. All experiments: bars and error bars represent mean ± SEM. \**p* < 0.05; \*\**p* < 0.01; \*\*\**p* < 0.001; ns, not significant.

### Increased ROS levels after LDHA inhibition are not responsible for the observed AML cell death

In a previous study in mice, it was shown that loss of LDHA in HSCs increases ROS levels because of increased OXPHOS activity, and these excessive ROS levels disrupt HSC maintenance [19]. However, AML cell death upon loss of LDHA could not be clearly linked to ROS. We therefore wondered whether LDHA inhibition in human AML cells would augment ROS levels due to increased OXPHOS activity, and could cause the observed cell death. We measured ROS levels in AML cell lines (n=8) by flow cytometry using a CellROX dye after exposing the cells to increasing concentrations of FX11 for 24 hours. ROS levels were significantly increased up to 5-fold as compared to control samples (Fig. 3A). Since the antioxidant N-acetyl-L-cysteine (NAC) has been reported to decrease ROS in HSCs [19], we performed a rescue experiment by supplementing NAC post-LDHA inhibition to understand if excess ROS contributes to AML cell death. Interestingly, although NAC supplementation reduced ROS nearly to control levels (Fig. 3B), cell viability was not rescued upon LDHA inhibition (Fig. 3C). As an independent approach, we also co-cultured AML cells with bone marrow derived mesenchymal stromal cells (MS-5 cell line) that were previously shown to reduce ROS levels in AML cells [20]. Similar to our above-mentioned results with NAC supplementation, MS-5 co-cultures significantly reduced ROS levels (Fig. 3D) but cell viability after LDHA inhibition was not rescued (Fig. 3E). Similar results were obtained with primary AML patient samples (Supplementary Fig. 1A). Collectively, these findings suggest that excessive ROS levels induced by LDHA inhibition are not the main cause of cell death in AML cells.

**Figure 3.**
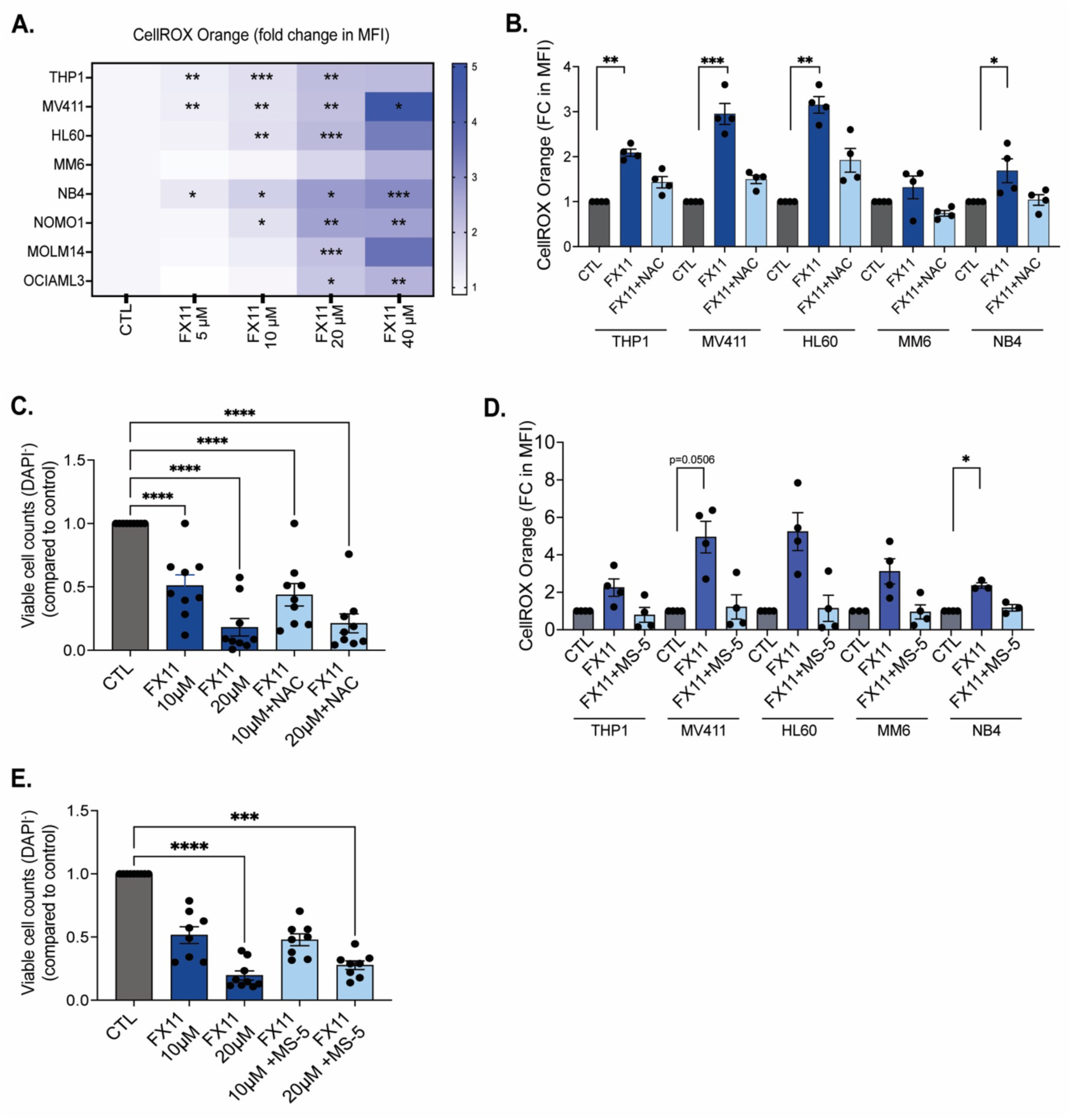
Increased ROS levels do not contribute to AML cell death following LDHA inhibition. **A.** Heatmap showing the fold change of CellROX Orange mean fluorescent intensity in AML cell lines exposed to increasing concentrations of FX11 for 24 hours, compared to DMSO treated controls. Data show mean of biological triplicates. **B**. Fold change (FC) of CellROX Orange mean fluorescent intensity (MFI) in AML cell lines exposed to 20 µM FX11 in the presence and absence of NAC (N-acetyl-L-cysteine, 2 mM) for 24 hours. Each dot represents a biological replicate measured in technical triplicate. **C**. Number of viable (DAPI^-^) cells relative to controls in AML cells (each dot represents a distinct AML cell line, as mean of biological replicates) following 20 µM FX11 treatment in the presence and absence of 2 mM NAC for 24 hours. **D**. Fold change of CellROX Orange mean fluorescent intensity in AML cell lines exposed to 20 µM FX11 in the presence (co-culture) and absence (mono-culture) of MS-5 cells for 24 hours. Each dot represents a biological replicate measured in technical triplicate. **E**. Number of viable (DAPI^-^) cells relative to controls in AML cells (each dot represents a distinct AML cell line, as mean of biological replicates) following 20 µM FX11 treatment in the presence and absence of MS-5 cells for 24 hours. **B**,**C,D,E**: one-way ANOVA; **A**: two-way ANOVA for multiple comparisons. All experiments: bars and error bars represent mean ± SEM. \**p* < 0.05; \*\**p* < 0.01; \*\*\**p* < 0.001; \*\*\*\**p* < 0.0001.

### Global metabolic changes induced in AML cells after LDHA inhibition

To further define the metabolic status of AML cells after LDHA inhibition, we measured intracellular metabolites after 15 minutes and 24 hours of treatment with FX11, assessing short-term and long-term consequences respectively. Metabolites were extracted from 3 different AML cell lines and analyzed by liquid chromatography-coupled mass spectrometry (LC-MS). After normalization of the samples to the internal standard (d8-valine), we investigated common changes occurring in the 3 AML cell lines across different metabolic pathways, including glycolysis, the pentose phosphate pathway (PPP), hexosamine biosynthesis, the TCA cycle and redox balance after 24 hours (Fig. 4A). Several upstream glycolytic metabolites such as hexose, glucose-6-phosphate and fructose-6-phosphate were significantly reduced after 15 minutes and 24 hours of treatment with FX11 (Fig. 4B) in AML cells. Expectedly, intracellular lactate levels were significantly reduced after FX11 treatment in NB4 cells (Fig. 4C), which was further confirmed across different AML cell lines (Supplementary Fig. 1B, n=3), showing reduced pyruvate fermentation to lactate.

**Figure 4.**
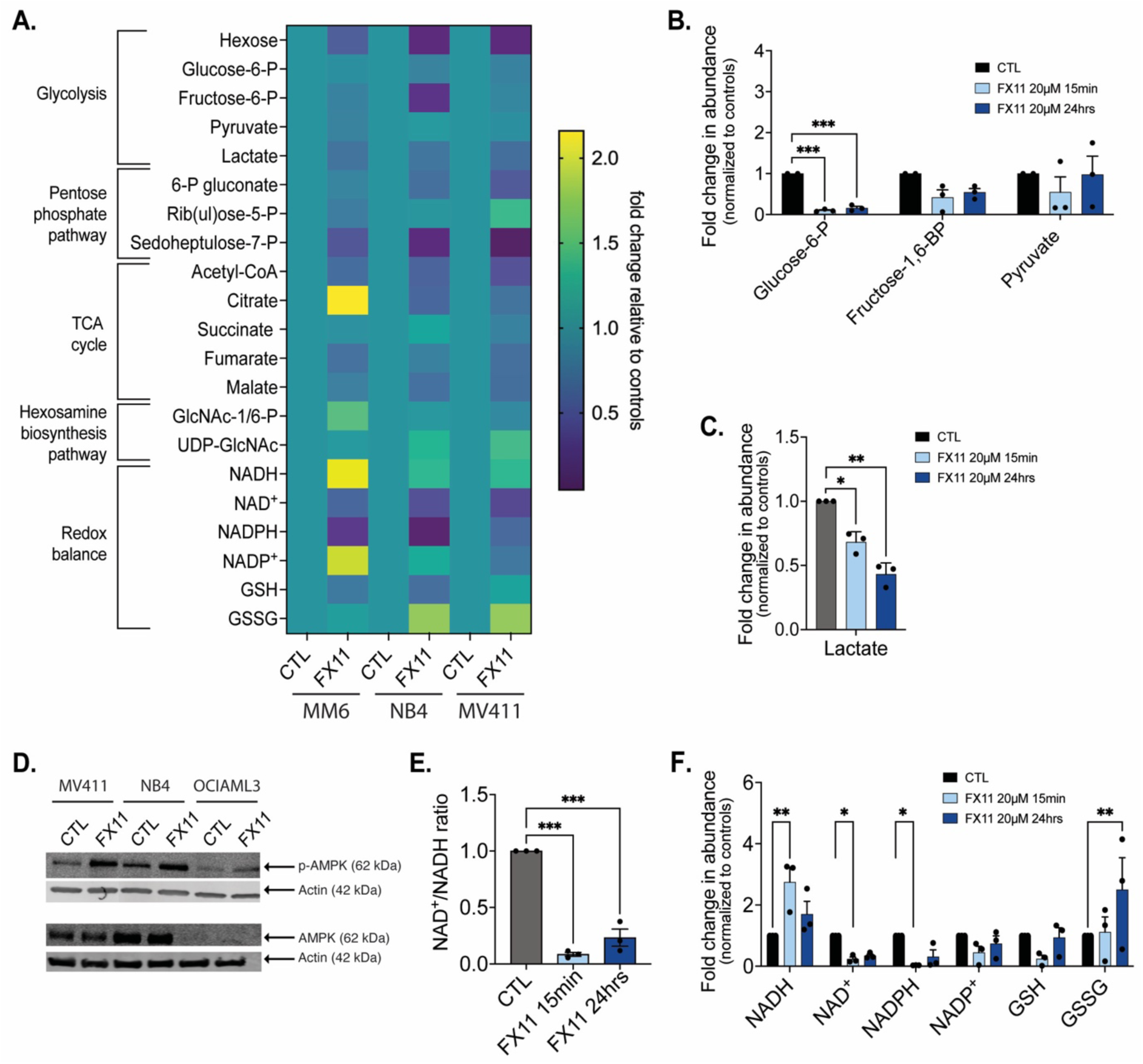
LDHA inhibition induces global metabolic changes. **A.** Heatmap showing metabolomics results of AML cells (n=3) after 24 hours of treatment with 20 µM FX11. Data show mean of technical triplicates as fold change compared to vehicle-treated groups. **B-C**. Glycolysis-related metabolites (B) and intracellular lactate abundance (C) in NB4 cells after 15 minutes and 24 hours of treatment with 20 µM FX11, as measured by LC-MS (each dot represents a biological replicate). **D**. Western blot for p-AMPK and AMPK in AML cells lines grown for 24 hours in the presence or absence of 20 µM FX11. β-actin was used as loading control. Image shown is a representative of three independent replicates**. E**. NAD^+^/NADH ratio in NB4 AML cells after 15 minutes and 24 hours of treatment with 20 µM FX11, as measured by LC-MS (each dot represents a biological replicate). **F**. Relative abundance of redox homeostasis related metabolites in NB4 cells after 15 minutes and 24 hours of treatment with 20 µM FX11, as measured by LC-MS (each dot represents a biological replicate). **B,C,E,F**: one-way ANOVA. All experiments: bars and error bars represent mean ± SEM. \**p* < 0.05; \*\**p* < 0.01; \*\*\**p* < 0.001.

Besides the effect on glycolysis, treatment of AML cells with FX11 for 24 hours reduced the levels of several PPP metabolites, while changes in hexosamine biosynthesis (Fig. 4A) and in amino acids (Supplementary Fig.1C) were less evident. When examining the levels of TCA cycle metabolites, we found an overall reduction across the different cell lines after FX11 treatment (Fig. 4A). The reason for this was unclear, given the increase in OCR in AML cells treated with LDHA inhibitors (Fig. 2E, F). To further examine this, we measured the mitochondrial membrane potential of AML cells (n=8) exposed to increasing doses of FX11, which confirmed higher OXPHOS activity (Supplementary Fig. 1D). Additional experiments revealed that this increase in membrane potential was associated with an increase in lipid uptake (Supplementary Fig. 1E), suggesting that AML cells may increase fatty acid oxidation in response to inhibition of glycolysis, resulting in higher OXPHOS activity.

Both glycolysis and OXPHOS can be used by cells for ATP generation to meet bioenergetic demands. To assess whether the observed increase in OXPHOS was sufficient to compensate for reduced glycolytic activity upon LDHA inhibition in AML cells, we measured phosphorylation of 5’ AMP-activated protein kinase (AMPK), which acts as a key cellular energy sensor. In all cell lines tested we detected increased AMPK phosphorylation in the presence of FX11 (Fig. 4D), indicating that even though OXPHOS activity increases, AML

cells critically depend on LDHA to maintain their energy balance. Additional supplementation of cells with different anaplerotic substrates, including pyruvate, glutamine or a cell-permeable form of 2-oxoglutarate, to increase the levels of TCA cycle intermediates could however not prevent the induction of cell death by LDHA inhibition (Supplementary Fig. 1F), showing that the lower levels of TCA cycle metabolites are not responsible for the observed cell death.

Given the importance of glycolysis and the PPP in regulating the cellular redox status, we next looked more carefully at the levels of NADH, NAD^+^, NADPH, NADP^+^, reduced glutathione (GSH), oxidized glutathione (GSSG) and their respective ratios. In line with the role of LDHA in regenerating NAD^+^ from NADH, we found increased levels of NADH, decreased levels of NAD^+^ and a decreased NAD^+^/NADH ratio in FX11-treated AML cells at both early and later time points (Fig. 4E, F). We also found decreased levels of NADPH relative to NADP^+^ and higher GSSG versus GSH levels (Fig. 4F), in line with a decreased PPP activity and increased demand for ROS detoxification (see Fig. 3). Since we already ruled out increased ROS as the reason for LDHA inhibition-induced AML cell death, we next focused on the role of LDHA in maintaining the cytosolic NAD^+^/NADH ratio.

### LDHA is essential to maintain the intracellular NAD^+^/NADH balance in AML cells

LDHA catalyzes the interconversion of pyruvate and lactate with concomitant interconversion of reduced NADH and NAD^+^. In glycolysis, LDHA ferments pyruvate to lactate and thereby regenerates NAD^+^, an essential co-factor for the central glycolytic enzyme glyceraldehyde 3-phosphate dehydrogenase (GAPDH). Given this role of LDHA and the stable decline in NAD^+^ levels in different AML cell lines post-LDHA inhibition (Fig. 5A), we decided to further study the consequences of the disturbance in intracellular NAD^+^/NADH balance. We first validated our metabolomics results with an independent approach by monitoring the cytosolic NAD^+^/NADH ratio using a genetically-encoded fluorescent biosensor. Peredox-mCherry was designed with a bacterial NADH-binding protein linked to a circularly permuted green fluorescent T-Sapphire and a NADH-insensitive red fluorescent mCherry, allowing measurement of the cytosolic free NAD^+^/NADH ratio as the red-to-green fluorescence ratio [21, 22]. We transduced NB4 cells (a glycolytic and LDHA inhibitor-sensitive AML cell line) with a Peredox-mCherry encoding lentiviral vector and sorted mCherry^+^ cells. When these cells were treated for 15 minutes with either FX11 or GSK2837808A, an increase in the green fluorescent signal (FITC channel) relative to mCherry signal was detected by flow cytometry (Fig. 5B), confirming that LDHA inhibition decreases the NAD^+^/NADH ratio.

**Figure 5.**
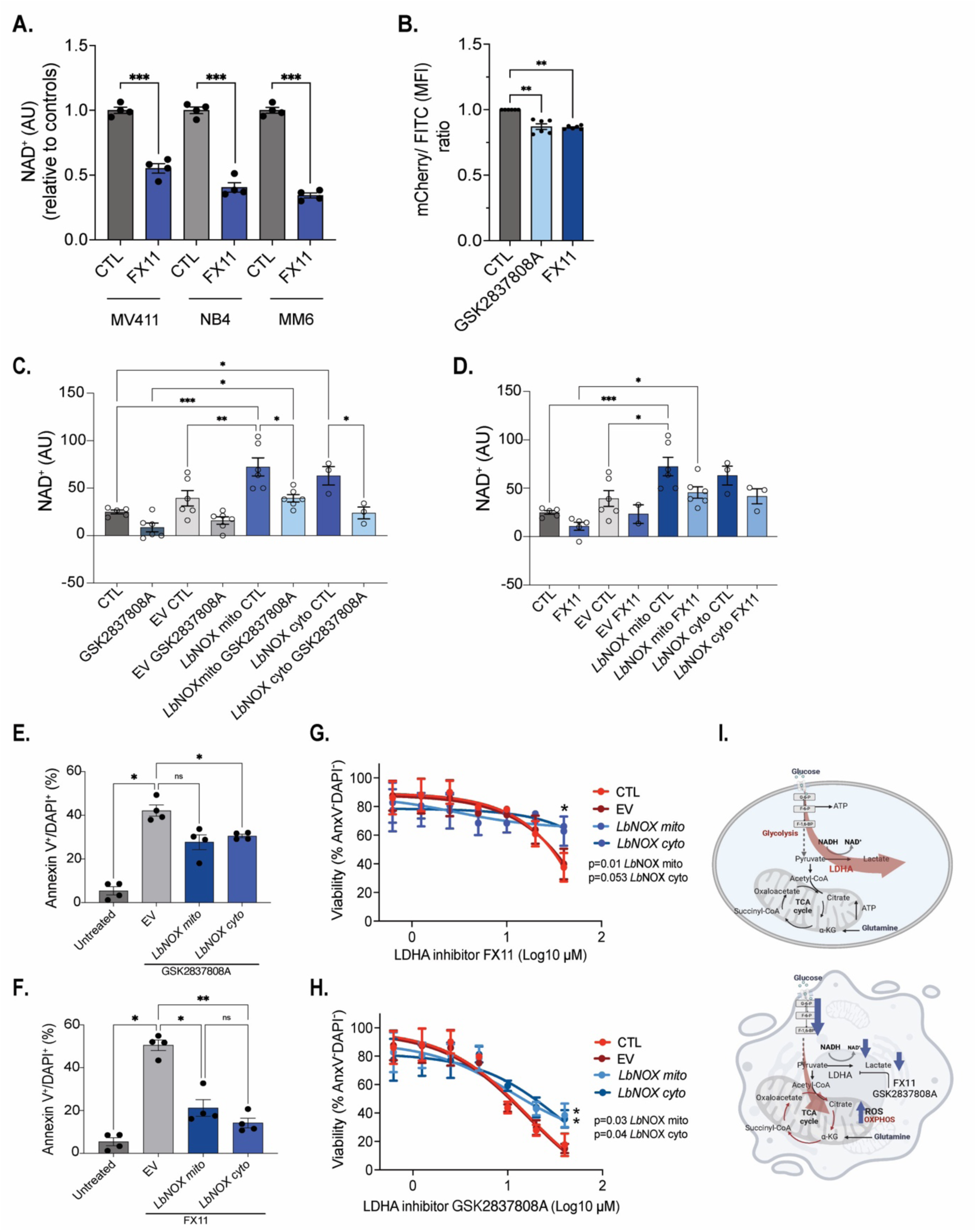
LDHA critically regulates the intracellular NAD^+^/NADH ratio in AML cells. **A.** Intracellular NAD^+^ abundance in AML cell lines after 24 hours of treatment with 20 µM FX11, as measured by LC-MS (n=4 technical replicates). **B**. Red (insensitive to NADH)-to-green (sensitive to NADH) fluorescence ratio in Peredox-mCherry-expressing NB4 cells treated for 15 minutes with either FX11 or GSK2837808A. Each dot represents a biological replicate. **C-D**. Cellular NAD^+^ levels in NB4 control cells, NB4 cells transduced with an empty vector (EV) and NB4 cells overexpressing a mitochondrial or cytosolic variant of *Lb*NOX, treated with vehicle or 20 µM GSK2837808A (C) or 20 µM FX11 (D) for 24 hours, as measured by enzymatic assay. **E-F**. Percentage dead (Annexin V^+^DAPI^+^) NB4 control cells, NB4 cells transduced with an EV and NB4 cells overexpressing a mitochondrial or cytosolic variant of *Lb*NOX, treated with vehicle or 20 µM GSK2837808A (E) or 20 µM FX11 (F) for 24 hours. **G-H**. Percentage live (Annexin V^-^DAPI^-^) NB4 control cells, NB4 cells transduced with an EV and NB4 cells overexpressing a mitochondrial or cytosolic variant of *Lb*NOX, treated with increasing doses of GSK2837808A (G) or FX11 (H) for 18 hours. **I**. Graphical abstract depicting the metabolic role of LDHA in AML cells (upper panel) and consequences of LDHA inhibition (lower panel). G-6-P = Glucose 6-phosphate, F-6-P = Fructose 6-phosphate, F-1,6-BP = Fructose 1,6-bisphosphate. **A-F**: one-way ANOVA; **G,H**: two-way ANOVA for multiple comparisons. All experiments: bars and error bars represent mean ± SEM. \**p* < 0.05; \*\**p* < 0.01; \*\*\**p* < 0.001; ns, not significant.

To test whether the reduced NAD^+^/NADH ratio was responsible for cell death induction upon LDHA inhibition in AML cells, we tested whether re-balancing the NAD^+^/NADH ratio could rescue AML cell death. To this end, we transduced NB4 cells with lentiviral constructs overexpressing a mitochondrial or cytosolic variant of the water-forming NADH oxidase (NOX) from *Lactobacillus brevis* (*Lb*NOX) [23]. Quantification of intracellular NAD^+^ levels using a colorimetric assay showed that stable expression of either mitochondria-or cytosol-targeted *Lb*NOX increased cellular NAD^+^ levels compared to control cells and normalized NAD^+^ levels in NB4 cells treated with GSK2837808A (Fig. 5C) or FX11 (Fig. 5D) which was also reflected in the NAD^+^/NADH ratios (Supplementary Fig.1 G, H). Analysis of AML cell death post-LDHA inhibition with GSK2837808A or FX11 showed that restoration of cellular NAD^+^ levels by *Lb*NOX significantly reduced cell death, although it was not fully prevented (Fig. 5E, F). In accordance, cell viability was significantly higher in *Lb*NOX-overexpressing cells compared to controls when treated with increasing doses of GSK2837808A or FX11 (Fig. 5G, H). Collectively, these findings show that LDHA plays a direct and central role in maintaining the NAD^+^/NADH balance in AML cells (Figure 5I, upper panel), and that the absence of this enzyme leads to a metabolic imbalance that cannot be compensated for by other enzymes, ultimately triggering cell death in AML cells (Figure 5I, lower panel).

## Discussion

AML blasts display a typical Warburg metabolism with high rates of aerobic glycolysis supporting the bioenergetic and biosynthetic demands of increased proliferation, redox regulation, hyperactive signaling cascades and therapy-induced damage repair [10, 13, 15, 19]. Unlike certain solid tumor types, where the role of individual glycolytic enzymes and their therapeutic potential has been studied in detail [24], a good strategy for targeting glycolytic metabolism in AML cells has remained elusive. In the current study, we showed that among glycolytic enzymes LDHA protein levels show the highest correlation with glycolytic activity in AML cells, and that LDHA inhibition shows potent anti-leukemic effects in both AML cell lines and primary patient samples, while sparing non-malignant hematopoietic cells. We further found that inhibition of LDHA impacts AML cell viability by lowering the cellular NAD^+^/NADH ratio, leading to reduced ATP generation through glycolysis that cannot be compensated for by increased mitochondrial OXPHOS activity.

One of the first studies revealing the dependence of AML blasts on aerobic glycolysis was that of Wang *et al*., where they showed that loss of either PKM2 or LDHA enforced a metabolic shift from glycolysis to mitochondrial respiration and strongly inhibited leukemia initiation and maintenance in mice [19]. Although the effects of PKM2 deletion on normal hematopoiesis were relatively mild *in vivo*, HSCs from *Ldha*^−/−^ mice exhibited increased ROS levels that negatively affected their functioning. However, depletion of LDHA did not impair the ROS buffering capacity of AML cells, suggesting distinct effects of LDHA loss on normal and malignant HSPCs. In line with their observation, we found that excessive ROS levels resulting from LDHA inhibition in human AML cells were not responsible for the observed cell death. Instead, in AML cells LDHA is key to maintain an optimal cytosolic NAD^+^/NADH ratio that allows maximal glycolytic flux and prevents cell death. In contrast to the Wang *et al*. study, where mouse HSCs, HPCs and AML cells were all shown to be equally impacted by complete loss of LDHA, we found that normal human HSPCs are less sensitive to LDHA inhibitors compared to AML blasts. As indicated by our extracellular flux (Fig. 2) and metabolomics (Fig. 4) analysis, the concentrations of LDHA inhibitors that we used in our study decrease but do not completely shut down LDHA activity, suggesting the existence of a therapeutic window, where partial inhibition of LDHA kills leukemic but not healthy HSPCs.

In the current study we focus on bulk AML cells, but recent findings from us and others shows that LSCs, or at least a subset of them, also depend on a high glycolytic flux. Using the genetically encoded metabolic sensor SoNar, which like Peredox reports the cytosolic NADH/NAD^+^ ratio, it was shown that cells with high cytosolic NADH/NAD^+^ ratios are highly glycolytic, enriched for LSCs, and reside in the hypoxic endosteal niche in an MLL-AF9-driven model of mouse AML [13]. In this model, pyruvate dehydrogenase kinase 2 (PDK2) was found to be essential for preventing pyruvate entry into the mitochondria and maintaining glycolytic activity and LSC properties. In our own previous work in human AML, we found that PDK1^high^ AMLs are highly glycolytic, have low OXPHOS and are enriched for stemness signatures [10]. In addition, these AMLs are mostly wildtype for FLT3 and NPM1, in contrast to FLT3-ITD and NPM1^c^ AMLs that are OXPHOS-dependent [10, 11]. It will be of interest to explore the therapeutic potential of LDHA inhibitors in future *in vivo* studies, for example using patient-derived AML xenografts harboring different genetic mutations, to test the efficacy of LDHA inhibitors in eradicating LSCs with high glycolytic signatures, investigate the effects in AMLs carrying metabolically-distinct subclones and study potential escape mechanisms.

An important finding of our study is that LDHA is absolutely essential for maintaining the optimal NAD^+^/NADH ratio in AML cells, and that this function cannot by compensated for by other endogenous NAD^+^ regenerating systems such as the malate-aspartate shuttle or the glycerol-3-phosphate shuttle that move glycolysis-derived NADH into the mitochondria for oxidation. Indeed, inhibition of LDHA increased the OCR and mitochondrial membrane potential while lowering the levels of TCA cycle metabolites, which may indicate that excessive shuttling of cytosolic NADH to the mitochondria for oxidation leads to inhibition of NAD^+^-dependent TCA cycle reactions. In accordance, both cytosolic and mitochondrial forms of *Lb*NOX rescued cellular NAD^+^/NADH ratios and viability, indicating that while NADH shuttling systems in AML cells are very active, increased activity of complex I of the electron transport chain cannot prevent a drop in the NAD^+^/NADH ratio when LDHA activity is reduced. Other NAD^+^ regenerating systems, such as NAD^+^ synthesis and salvage pathways, also do not seem to be able to compensate for the loss of LDHA. Tumor cells have been shown to increase NAD^+^ levels and upregulate NAD^+^ synthesis and salvage enzyme expression to support redox homeostasis and to fulfil the high demands of proliferation [25]. It has been recently noted that depleting NAD^+^ by inhibiting nicotinamide phosphoribosyltransferase (NAMPT), a key enzyme in the NAD^+^ salvage pathway, offers a therapeutical window in AML [26]. In LDHA-dependent lymphoma models, combinatory treatment of NAMPT inhibition and the LDHA inhibitor FX11 induced tumor regression in *in vivo* xenograft models [27]. In addition, a recent study highlighted that dual inhibition of the NADPH oxidase NOX2 and glycolysis — by targeting hexokinase or LDH — significantly reduced cell proliferation and delayed leukemia progression [28]. These findings, together with our current results, underscore the importance of redox metabolism to meet the particular bioenergetic, biosynthetic and antioxidant demands of AML cells. With this in mind, future studies should aim to further dissect the mechanisms by which genetically-distinct AMLs meet their redox demands and how this metabolic rheostat is impacted by different therapies. In addition, it will be worth to investigate if and how AML blasts and LSCs can adapt to different therapies impacting NAD^+^ pools or the NAD^+^/NADH balance. In this context, targeting LDHA holds promise in AML as it hits the Warburg effect at its core, shows a therapeutic window compared to healthy HSPCs and cannot be prevented by the presence of bone marrow stromal cells, which often protect AML cells from therapies.

## Methods

### Ex vivo AML and healthy primary cell cultures and *in vitro* cell culture conditions

Human AML cell lines (THP1, HL60, NB4, Mono-Mac-6 [MM6], MV411, NOMO1, MOLM14, OCIAML3 and U937, all from DSMZ) used in this study were maintained in RPMI-1640 medium (Gibco) supplemented with 10% fetal bovine serum (FBS, Sigma-Aldrich), 2 mM glutamine (ThermoFisher Scientific) and 1% antibiotics mix composed of penicillin and streptomycin (P/S; ThermoFisher Scientific). Cell line identity was validated by DNA fingerprinting. CD34^+^ human cord blood-derived cells were obtained from STEMCELL Technologies and were maintained in StemSpan SFEM-II with CD34 supplements (STEMCELL Technologies).

Primary AML specimens (CD34^+^ cells and mononuclear cells for CD34^-^ samples) were obtained after written informed consent at the Montefiore Medical Center/Montefiore Einstein Comprehensive Cancer Center (for patient characteristics see Supplementary Table 1). Frozen bone marrow aspirates were thawed in a water bath at 37 °C and resuspended in alpha-Minimum Essential Medium (α-MEM, ThermoFisher Scientific) supplemented with 2% FBS, DNase I (20 Units/mL), 4 mM MgSO_4_ and heparin (5 Units/mL) and incubated at 37 °C for 15 minutes. Primary AML cells were plated in Gartner’s medium consisting of α-MEM supplemented with 12.5% heat-inactivated FBS (Sigma-Aldrich), 12.5% heat-inactivated horse serum (Invitrogen), 1% P/S, 57.2 µM β-mercaptoethanol (Sigma-Aldrich) and 1 mM hydrocortisone (Sigma-Aldrich), with the addition of G-CSF (Amgen), N-Plate (Clinical grade TPO; Amgen) and IL-3 (Sandoz) (all 20 ng/mL). For AML cell line co-cultures, mouse bone marrow derived MS-5 stromal cells (DSMZ) were expanded in α-MEM with GlutaMAX, 10% FBS and 1% P/S, plated in culture flasks and expanded to form a confluent layer, to which AML cells were added. Liquid cultures and co-cultures were grown at 37 °C and 5% CO_2_.

We used HEK-293T cells to produce lentiviral particles, which were cultured in Dulbecco’s Modified Eagle’s Medium (D-MEM) (high glucose, ThermoFisher Scientific) supplemented with 10% FBS, 2 mM glutamine and 1% P/S.

### Reagents

Reagents purchased include: FX11 (HY-16214, Bio-connect), GSK2837808A (HY-100681, Bio-connect), AZD3965 (HY-12750, Bio-connect), Compound 3k (HY-103617, Bio-connect), N-acetyl-L-cysteine (NAC; A9165, Sigma-Aldrich). For flow cytometric reagents see section below, and Seahorse reagents are indicated in Seahorse assay section.

### Flow cytometry and sorting procedures

Cell death was quantified by Annexin V staining (FITC) according to manufacturer’s protocol (#560931, BD Biosciences). Equal number of cells were stained with Annexin V in calcium-supplied sterile water buffer for 20 minutes at 4 °C in the dark followed by DAPI staining for 10 minutes. CellROX Orange (C10443, ThermoFisher Scientific) was used at 1 µM final concentration in phosphate-buffered saline (PBS) for 15 minutes in the dark at 37°C. BODIPY-FL-C_16_ (4,4-Difluoro-5,7-Dimethyl-4-Bora-3a,4a-Diaza-*s*-Indacene-3-Hexadecanoic Acid, D3821, ThermoFisher Scientific; a green fluorescent fatty acid used to measure lipid uptake) was used at 1 µM final concentration in PBS for 15 minutes in the dark at 37 °C. MitoTracker Deep Red (M46753, ThermoFisher Scientific) was used at 500 nM final concentration in PBS for 15 minutes in the dark at 37°C. Analysis was performed on a FACSVerse system (BD Biosciences).

For monitoring the cytosolic NADH / NAD^+^ ratio, NB4 cells were lentivirally transduced with the NADH-sensitive Peredox probe under the control of a phosphoglycerate kinase (PGK) promoter [21]. After transduction, mCherry^+^ cells were sorted (FACSAria II; BD Biosciences). NADH-sensitive green fluorescence (FITC channel; mean fluorescence intensity) was recorded by flow cytometry (FACSVerse; BD Biosciences) and expressed relative to stable mCherry mean fluorescence intensity, before and after the treatment with FX11 for 15 minutes. The mCherry/FITC ratio therefore indicates changes in NAD^+^/NADH ratio. Results were analyzed using the FlowJo Software (BD Biosciences).

### Liquid Chromatography-coupled Mass Spectrometry (LC-MS)

1x10^6^ AML cells were collected after 24 hours of treatment with 20 µM FX11 and washed with sterile UltraPure DNase/RNase-Free Distilled Water (10977049, Life Technologies Europe). Cells were maintained on ice during the process to keep their metabolism stabilized. For the extraction, 200 µL of 9:1 Methanol/Chloroform (#1.06035 and #650471, Sigma-Aldrich), supplemented with 5 µM D8-valine (DLM-311-1, Buchem BV; internal standard) was added to the cells. Samples were vortexed and centrifuged at 13 000 rpm at 4 °C for 10 minutes. Supernatant was harvested and dried down with a SpeedVac machine for 2 hours approximately. Samples were stored at -80 °C until the LC-MS analysis was conducted. Right before the analysis, 45 µL of 80% methanol in water was added to the samples. From that, 30 µL was loaded in an LC-MS vial and untargeted LC-MS analysis was performed as previously described [29]. Briefly, 5 µL of sample was injected and separated on an Inertsil 3 µm particle ODS-4 column (150 x 2.1 mm; GL Biosciences) at a constant flow rate of 0.2 mL/min with an Agilent 1290 HPLC system. Mobile phase A consisted of 5 mM hexylamine (Sigma-Aldrich) adjusted to pH 6.3 with acetic acid (Biosolve BV) and phase B of 90% methanol (Biosolve BV)/10% 10 mM ammonium acetate (Biosolve BV) adjusted to pH 8.5 with ammonia (Merck). The mobile phase profile consisted of the following steps and linear gradients: 0 – 2 min at 0% B; 2 – 6 min from 0 to 20% B; 6 – 17 min from 20 to 31% B; 17 – 36 min from 31 to 60% B; 36 – 41 min from 60 to 100% B; 41 – 51 min at 100% B; 51 – 53 min from 100 to 0% B; 53 – 60 min at 0% B. Analytes were identified with an Agilent 6550 ion funnel mass spectrometer operated in negative mode with an electrospray ionization (ESI) source and the following settings: ESI spray voltage 3500 V, sheath gas 350 °C at 11 L/min, nebulizer pressure 35 psig and drying gas 200 °C at 14 L/min. An m/z range from 70 to 1200 was acquired with a frequency of 1 per second by adding 8122 transients. Compound identification was based on their exact mass (<5 ppm) and retention time compared to standards. The areas under the curve of extracted-ion chromatograms of the [M-H]^-^ forms were integrated using MassHunter software (Agilent, Santa Clara, CA, USA), and normalized to the mean of the areas obtained for a series of 150 other metabolites (‘total ion current’). Metabolites were further characterized in MS2 by fragmentation with 10, 20 and 40 V and 150 ms acquisition time per spectrum and a 4 mass unit isolation window.

### Seahorse assay

Oxygen consumption rate (OCR) and Extra Cellular Acidification Rate (ECAR) were measured using a Seahorse XF96 analyzer (Seahorse Bioscience, Agilent, US) at 37 °C. For AML cell lines 100.000 cells and for primary AML samples 200.000 cells were seeded per well in poly-L-lysine (Sigma-Aldrich) coated Seahorse XF96 plates in 180 µL XF Assay Medium (Modified DMEM, Seahorse Bioscience). For OCR measurements, XF Assay Medium was supplemented with 10 mM Glucose and 20 µM of FX11 or 10 µM of GSK2837808A (Port A), 2.5 µM oligomycin A (Port B), 2.5 µM FCCP (carbonyl cyanide-4-[trifluorometh oxy] phenylhydrazone) (Port C) and 2 µM antimycin A together with 2 µM Rotenone (Port D) were sequentially injected to measure basal and maximal OCR levels (all reagents from Sigma-Aldrich). For ECAR measurements, Glucose-free XF Assay medium was added to the cells and 20 µM of FX11 or 10 µM of GSK2837808A (Port A), 10 mM Glucose (Port B), 2.5 µM oligomycin A (Port C) and 100 mM 2-deoxy-D-glucose (Port D) were sequentially injected (all reagents from Sigma-Aldrich). The XF96 protocol consisted of 4 times mix (2 minutes) and measurement (2 minutes) cycles, allowing for determination of OCR/ECAR at baseline and in between injections. Both basal and maximal OCR levels as well as basal ECAR and glycolytic capacity measurements were calculated by assessing metabolic response of the cells in accordance with the manufacturer’s instructions. The OCR and ECAR measurements were normalized to the number of cells used for the assay. Seahorse Wave Desktop software (Agilent) was used to design assays and for data analysis.

### Lentiviral vectors and transduction

Cytosol-directed or mitochondria-directed *Lb*NOX (*Lactobacillus brevis* NADH oxidase) lentiviral constructs were provided by Dr. Jakub Rohlena (Czech Academy of Sciences) and were made by PCR amplifying *Lb*NOX or mito*-Lb*NOX cDNAs from the original pUC57 vectors (#75285, #74448, Addgene), cleaving the cDNAs with fast digest EcoRI and BamHI restriction enzymes(ThermoFisher Scientific) and ligated them into EcoRI and BamHI-cleaved pCDH-CMV-MCS-EF-puro lentiviral plasmids. Correct insertion was verified by sequencing. Lentiviruses were generated in HEK293T cells using empty vectors, cyto-*Lb*NOX or mito-*Lb*NOX (4 µg plasmid) together with 3 µg of delta 8.9 packaging plasmid and 2 µg of vesicular stomatitis virus (VSV-G) envelope plasmid using the Fugene transfection system (Promega, Madison, USA). After 48 hours, viral supernatant was harvested, filtered through a 0.45 µm filter, and used to transduce recipient cells (with 8 µg/mL polybrene to increase the infection efficiency). Cells were cultured in the presence of 1 µg puromycin for selection after transduction. The lentiviral vector containing the peredox probe was previously generated [21]. Briefly, the pcDNA3.1-Peredox-mCherry plasmid was obtained from the Addgene collection (plasmid #32383) and Peredox-mCherry was subcloned using a Gateway strategy into pTM978 (*PGK* > Gateway-IRES-Neo). Lentiviruses were generated as above.

### NAD^+^/NADH assay

Intracellular NAD^+^/NADH ratios were measured using a NAD^+^/NADH Quantitation kit (Sigma-Aldrich, MAK460) which allows to measure both total NAD^+^ and NADH levels in the cells. 1 × 10^6^ NB4 cells were seeded in culture plates for each condition, and the assay was performed after 24 hours of incubation with either 20 µM FX11 or GSK. We used equal protein amount (20 µg) of each sample (in technical duplicates or triplicates) in 96 well plates and recorded fluorescence using a GloMax® Discover spectrophotometer (Promega). We then calculated NAD^+^ and NADH levels using a standard curve and following the manufacturer’s instructions.

### Western blot

Cell pellets from AML cell lines (1 × 10^6^ cells) were lysed in RIPA buffer (#89900, ThermoFisher Scientific) supplemented with protease inhibitors (cOmplete Mini; #11836153001, Sigma-Aldrich) and phosphatase inhibitors (PhosSTOP, #4906837001, Sigma-Aldrich), and were sonicated in an ice bath at 25 kHz, power 100 for 1 minute. Lysates were then boiled at 100 °C for 5 minutes in 4x Laemmli sample Buffer (#1610747, Bio-Rad) containing 10% β-mercaptoethanol (M3148, Sigma-Aldrich). We loaded 25 µg of proteins/well into 4–15% mini–Protean TGX precast gels for SDS-PAGE (#4561085, Bio-Rad) and transferred proteins to 0.45 µm low fluorescence PVDF membranes (Trans-Blot Turbo RTA Mini Transfer kit, #1704274, Bio-Rad). Membranes were blocked in 5% BSA (A9418, Sigma-Aldrich) in Tris-buffered saline (TBS) 1X and incubated for 1 hour at room temperature on a rotary shaker. Then, membranes were incubated overnight with primary antibodies prepared in 5% BSA in TBS at 1:1000 dilution (rabbit anti-AMPK [2532S, Cell Signaling]: rabbit anti-phosphoAMPK [2535S, Cell Signaling]) + 1:50000 of mouse anti-actin (A5441, Sigma-Aldrich) used as a loading control. After several washes with TBS 1x + Tween 0.1% (TBST) (P1379, Sigma-Aldrich), secondary antibodies were added, prepared in 5% milk in TBS 1X; 1:30000 for actin (goat anti-mouse secondary antibody coupled with Dylight680; #35518, ThermoFisher Scientific) and 1:15000 for the test antibodies (goat anti-rabbit secondary antibody coupled with Dylight800, #SA-35571, ThermoFisher Scientific) . The membrane was placed on the shaker, in the dark, at room temperature for 1 hour. After, we washed the membranes 3 times for 5 minutes with TBST and 2 times for 5 minutes with TBS. An Odyssey XF Imager (LICORbio) was used to scan and analyze the membranes.

### Statistical analysis

All statistical analysis was performed using the GraphPad Prism Software. The means were plotted with errors bars representing standard error of the mean (SEM) with a minimal of 3 technical replicates and at least 3 biological replicates, unless otherwise indicated in the figure legend. Two-tailed student’s t-test were used to calculate the statistical significance between two groups, one-way ANOVA was used to calculate the statistical significance between three groups or more with one experimental parameter and two-way ANOVA for two or more experimental parameters. We chose an interval of confidence at 95% and the p-value was considered statistically significant if inferior to 0.05. For the figures, * = p-value < 0.05, ** = p-value < 0.01 and *** = p-value < 0.001.

## Acknowledgements

We thank Dr. Jakub Rohlena and Dr. Katherina Rohlenová from the Czech Academy of Sciences (Prague, Czech Republic) for kindly sharing the cytosol-directed and mitochondria-directed *Lb*NOX lentiviral constructs, and Dr. Amit Verma from the Albert Einstein College of Medicine (Bronx, NY, USA) for providing AML samples. We thank Dr. Nicolas Dauguet (de Duve Institute, UClouvain, Belgium) for the assistance with cell sorting. Schematics were created with BioRender.com.

## Supplementary Data

**Supplementary Figure 1.**
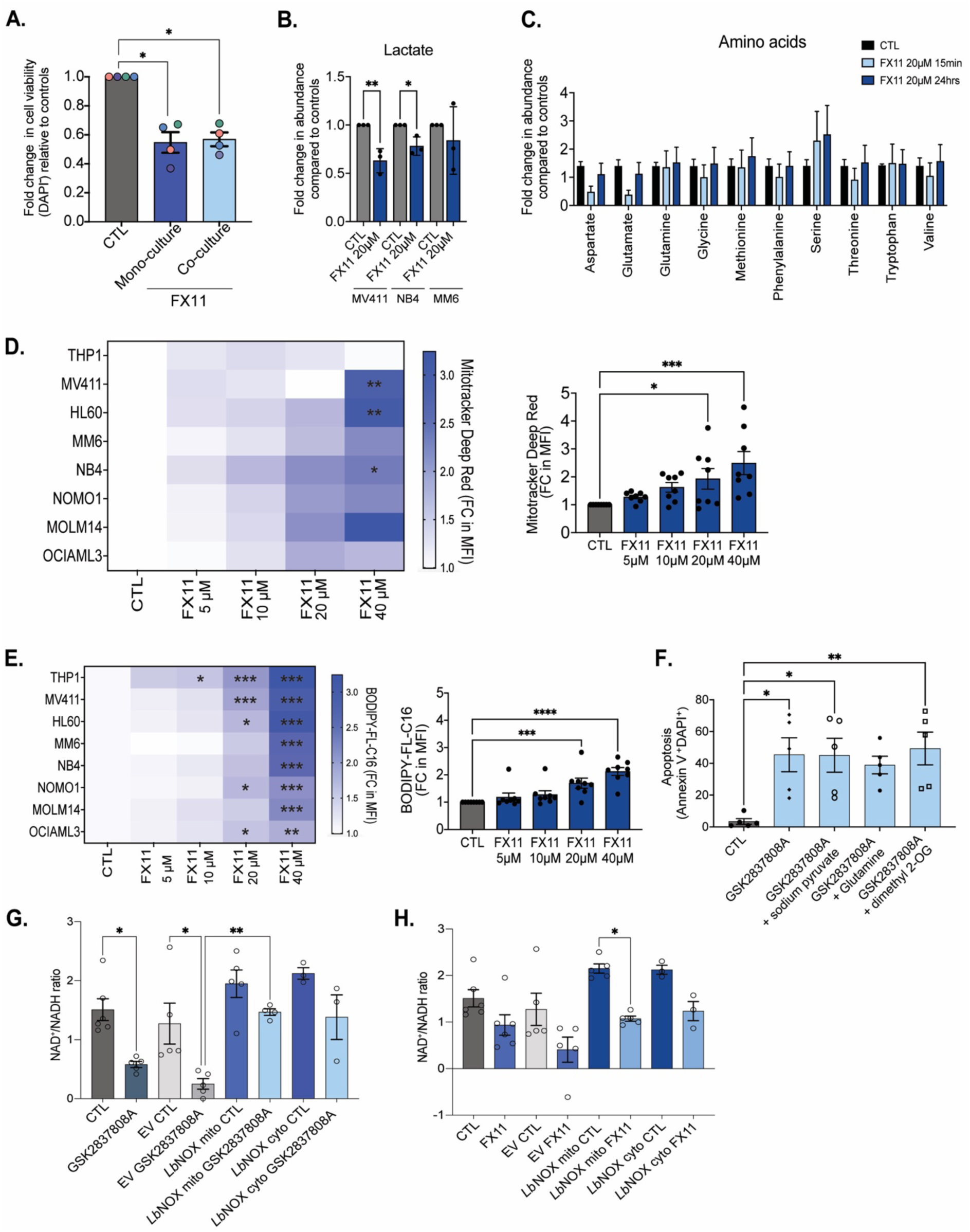
**A.** Number of viable (DAPI^-^) cells relative to controls in primary AML patient samples following 20 µM FX11 treatment in the presence (co-culture) and absence (mono-culture) of MS-5 cells for 48 hours. Each dot represents a distinct AML sample, as mean of biological triplicates. **B**. Intracellular lactate levels in different AML cell lines after 24 hours of treatment with 20 µM FX11, as measured by LC-MS. Each dot represents a technical replicate. **C**. Relative abundance of amino acids in NB4 cells after 15 minutes and 24 hours of treatment with 20 µM FX11, as measured by LC-MS. Bars show mean ± SEM of biological triplicates. **D**. Heatmap (left panel, per AML cell line tested, data show mean of biological triplicates) and bar graph (right panel, each dot shows a distinct AML cell line) showing the fold change of MitoTracker Deep Red mean fluorescent intensity in AML cell lines exposed to increasing concentrations of FX11 for 24 hours, compared to DMSO treated controls. **E**. Heatmap (left panel, per AML cell line tested, data show mean of biological triplicates) and bar graph (right panel, each dot shows a distinct AML cell line) showing the fold change of BODIPY-FL-C_16_ mean fluorescent intensity in AML cell lines exposed to increasing concentrations of FX11 for 24 hours, compared to DMSO treated controls. **F**. Percentage dead (Annexin V^+^DAPI^+^) AML cells treated with 20 µM GSK2837808A alone or with 2 mM sodium pyruvate, 2 mM glutamine or 400 µM dimethyl 2-OG (2-oxoglutarate) for 24 hours. Each dot represents a distinct AML cell line, as mean of biological triplicates. **G-H**. Cellular NAD^+^/NADH ratio in NB4 control cells, NB4 cells transduced with an empty vector (EV) and NB4 cells overexpressing a mitochondrial or cytosolic variant of *Lb*NOX, treated with vehicle or 20 µM GSK2837808A (G) or 20 µM FX11 (H) for 24 hours, as measured by enzymatic assay. **B**: Student’s t test; **A,D,E,F,G,H**: one-way ANOVA for multiple comparisons. All experiments: bars and error bars represent mean ± SEM. \**p* < 0.05; \*\**p* < 0.01; \*\*\**p* < 0.001; \*\*\*\**p* < 0.0001.

**Supplementary Table 1:** Patient characteristics Sample ID Sample information & known mutations.

| **Sample ID** | **Sample information & known mutations** |
| --- | --- |
| AML1 | BCORL1, DNMT3A, NRAS, PTPN11, TET2, U2AF1I |
| AML2 | FLT3-ITD, NPM1, DNMT3A, TP53 |
| AML3 | MLL rearrangement |
| AML4 | BCOR, DNMT3A, NRAS, RUNX1, STAG2 |
| AML5 | AML (no CD34<sup>+</sup> cells) |
| AML6 | KIT, PTPN11, WT1 |
| AML7 | NPM1, TET2 |
| AML8 | ASXL1 germline |
| AML9 | Chronic neutrophilic leukemia (CNL) with classic activating mutation in CSF3R (CSF3R-T618I). |
| AML10 | Venetoclax resistant AML; FLT3, IDH2, RUNX1, TET2 |
| AML11 | FLT3-ITD, NPM1, WT1 frameshift |
| AML12 | DNMT3A, NPM1, WT1 |
| AML13 | BRAF, GNAS, JAK2, NRAS, TET2 |
| AML14 | DNMT3A, FLT3, NPM1 |
| AML15 | CEBPA |

